# Genetic variation in trophic avoidance shows fruit flies are generally attracted to bacterial pathogens

**DOI:** 10.1101/2024.05.09.593162

**Authors:** Katy M. Monteith, Phoebe Thornhill, Pedro F. Vale

## Abstract

Pathogen avoidance behaviours are often assumed to be an adaptive host defence from infection. However, there is limited experimental data on the prerequisite for this assumption: heritable, intrapopulation phenotypic variation for avoidance. We investigated trophic pathogen avoidance in 122 inbred *Drosophila melanogaster* lines, and in a derived outbred population. Using the FlyPAD system, we tracked the feeding choice that flies made between substrates that were either clean or contained a bacterial pathogen. We uncovered significant, but weakly heritable variation in the preference index among fly lines. However, instead of avoidance, most lines demonstrated a preference for several bacterial pathogens, showing avoidance only for extremely high bacterial concentrations. Bacterial preference was not associated with susceptibility to infection and was retained in flies with disrupted immune signalling. Phenotype-genotype association analysis indicated several novel genes (*CG2321, CG2006, ptp99A*) associated with increased preference for the bacterial substrate, while the amino-acid transporter *sobremesa* was associated with greater aversion. Together with previous work showing fitness benefits of consuming high-protein diets, our results suggest that bacterial attraction may instead reflect a dietary preference for protein over carbohydrate. More work quantifying intrapopulation variation in avoidance behaviours is needed to fully assess its importance in host-pathogen evolutionary ecology.

## Background

Behavioural immunity is an animal’s first line of defence against infection and is characterized by behaviours that help avoid contact with infectious environments or infected conspecifics [1–4]. Invertebrates have evolved sophisticated mechanisms to detect and avoid pathogens involving the peripheral nervous system, specifically the gustatory and olfactory systems [5–11]. For example, geosmin, a microbial odorant, activates a specific subclass of olfactory neurons and inhibits oviposition, chemotaxis, and feeding, inducing pathogen avoidance [11]. More direct detection of pathogens also occurs via the gustatory sensory system of Drosophila, which detects bacterial cell wall components like lipopolysaccharide (LPS) and peptidoglycan (PGN). LPS is detected by bitter gustatory receptor neurons in the fly oesophagus expressing *TrpA1*, triggering feeding and oviposition avoidance [12,13], whereas PGN triggers grooming behaviour on stimulation of wing margins and legs [14] and learned avoidance of bacteria during feeding [15].

In contrast to the great detail learned about the mechanistic basis of avoidance behaviours in a few model systems [9], we currently know little about the extent of intrapopulation phenotypic or genetic variation in pathogen avoidance behaviours, the extent to which such variation is heritable, and therefore if it is likely to respond to selection [2]. Investigating the genetic and environmental factors that contribute to within-population variation in avoidance behaviours is important to understand its ecological and evolutionary consequences [2,16,17]. For instance, foraging and feeding are essential components of host ecology and are crucial for organismal reproduction and fitness, but they also provide a major route of pathogen transmission [18–22]. Characterising variability in host behaviours that contribute to individual heterogeneity in pathogen acquisition and spread is therefore a major focus of disease ecology and epidemiology [17,18,23–25].

Intrapopulation variation in avoidance behaviours is also likely to have evolutionary consequences for both hosts and pathogens [2,26]. The ability to avoid contact with pathogens allows healthy individuals to evade the pathology resulting from infection and further prevents the activation of the immune response, which may be metabolically costly or even cause immunopathology, with detrimental effects on host fitness [27–29]. Given that individuals vary widely in susceptibility and in the extent to which they experience losses in fitness during infection [30–33], the potential costs and benefits of avoiding infection are therefore likely to vary between individuals [2,17]. Such variable costs, in turn, may contribute to the maintenance of standing genetic variation in avoidance via fluctuating selection [34–36], and affect the predicted evolutionary trajectories of pathogens [37,38]. However, without empirical measurements of the phenotypic and genetic variation in pathogen avoidance behaviours, it is difficult to assess their true impact on host-pathogen ecology and evolution[2,26].

In the present work, we aimed to quantify the intrapopulation variation in behavioural pathogen avoidance in *Drosophila melanogaster*, a widely used model organism for host-pathogen interactions and behavioural genetics[39–41]. We designed a two-choice feeding assay using the FlyPAD system for tracking, recording, and analysing the feeding behaviour of fruit flies in real-time [42]. We assayed the preference for (or aversion of) the bacterial pathogen *Pseudomonas entomophila* in a panel of 122 inbred DGRP lines and avoidance of several bacterial pathogens in a DGRP-derived outbred population of flies. Our assay offered flies a choice of feeding from a clean substrate or an identical substrate to which an exponentially growing inoculum of bacteria had been added. This assay therefore mimicked a frequent natural scenario where insects must make foraging decisions between food sources that may or may not contain sources of pathogenic infection [19,21,22,43].

## Results

### Variation in fly aversion to a bacterial feeding substrate

Fruit flies are naturally attracted to bacterial volatiles [44–47], but previous work has also established that adult *D. melanogaster* actively avoid sources of pathogenic bacterial infection [13,15,48]. Our starting point was therefore to investigate the extent to which pathogen avoidance varied within a population, and how much of this variation could be explained by all sources of genetic variation, that is, the broad sense heritability *H*^*2*^ [49,50].

We exposed 122 DGRP lines to a choice between a clean substrate of 5% sucrose and an identical substrate to which an overnight culture (OD_600_= 1) of the bacterial pathogen *Pseudomonas entomophila* was added. We observed substantial among line genetic variation (Line effect p <0.001), with mean preference index scores ranging from to 0.22 (SE ± 0.16 – RAL-380) indicating pathogen avoidance, to −0.66 ± 0.06 (RAL-879) indicating strong attraction to food contaminated with *P. entomophila*. Contrary to our initial expectation, we found that the majority of DGRP lines (108 / 122) showed a preference for the pathogen-contaminated substrate, with a mean preference index across all 122 lines of −0.22 (median −0.31) (**Figure 1A**, **Table 1**). We detected a non-zero, but weak, genetic component underlying this variation in preference, with broad sense heritability estimate of 0.07-0.08 (Table 1).

**Table 1.** Descriptive statistics and variance components of fly preference index. We calculated the broad-sense heritability by partitioning the variance into genetic variance and residual variance caused by other factors. We calculated broad-sense heritability as *H*^2^ = *V*_*G*_/(*V*_*G*_ + *V*_*E*_) in two separate but similar models. In one analysis we fit a model with a binomial error structure, where the response variable included the total number of sips taken by flies on each substrate; in a second model, we modelled the preference index in a linear model. Both approaches yielded very similar estimates of variance components.

| Variance component | Binomial model using the number of sips on each substrate as the response | Linear model with preference index as the response |
| --- | --- | --- |
| *Mean* | -0.22 | - |
| *Median* | -0.31 | - |
| *Genetic Variance ($V_G$)* | 0.25 | 0.25 |
| *Phenotypic Variance ($V_P$)* | 3.53 | 3.29 |
| *Environmental Variance ($V_E$)* | 3.28 | 3.04 |
| *Broad-sense Heritability $H^2$* | 0.071 | 0.077 |

**Figure 1.**
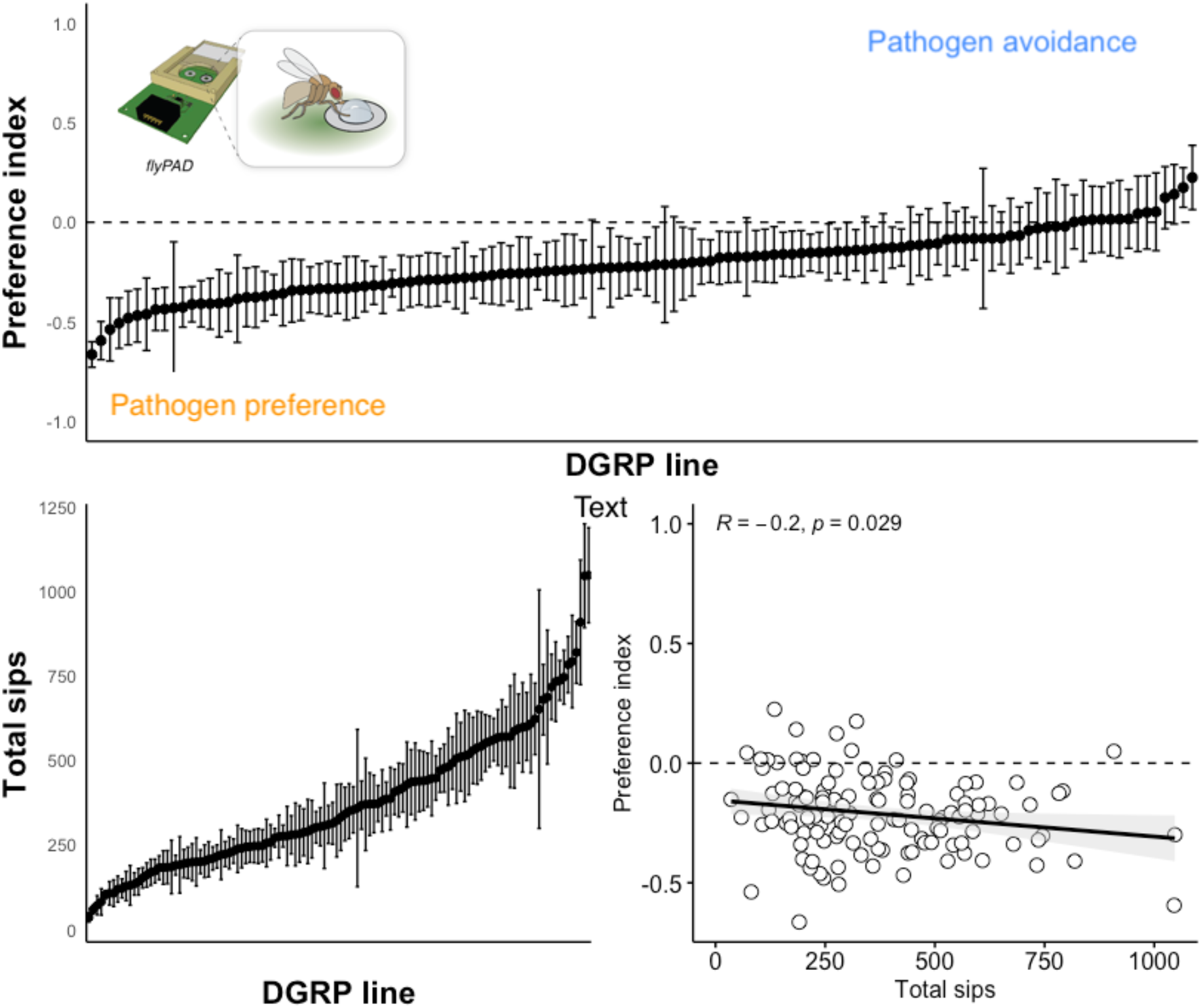
Genetic variation. **A**. We measured the choice index in 122 lines from the Drosophila Genetic Reference Panel (DGRP). We measured pathogen avoidance as a preference index, calculated over 30 minutes, as *(sips*_*clean*_ *-sips*_*pathogen*_*) / sips* _*total*_). Positive values of the preference index indicate pathogen avoidance, where a value of 1 indicates complete preference for the clean substrate; negative values suggest a pathogen preference, where −1 indicates complete preference for the pathogen-contaminated substrate; 0 indicates an equal number of sips on each substrate. Each line was replicated between 16-24 times. B) Line means for the total number of sips taken during a 30-min FlyPAD feeding assay. C. The relationship between the preference index in A and the total number of sips shown in B. Each data point is the line mean for each trait. R is the Pearson correlation coefficient, and the line shows the linear relationship, shaded by the 95% CI. FlyPAD image from https://flypad.rocks/.

Using the same feeding data, we were also able to calculate the total number of sips taken on both substrates by each fly line, as a measure of feeding activity. Feeding activity is relevant in the context of pathogen exposure because more active feeders may be more likely to acquire infection orally if they do not avoid contaminated substrates[51]. This analysis also revealed substantial genetic variation in feeding (Line effect p <0.001) ranging from a mean of 35.5±14 sips (RAL-790) to 1047±140 sips (RAL 059) measured over a 30-minute period (**Figure 1B**). These results are consistent with previous measures of genetic variation in food intake in the DGRP panel [52]. We identified a significant (but weak) negative relationship between the number of sips and the preference index (R= −0.2, p=0.029), indicating that fly lines with more active feeding also showed a stronger preference for the pathogen-contaminated substrate – although many low feeders also showed a preference for the pathogen-contaminated substrate (**Figure 1C**).

### Drosophila are attracted to several bacterial pathogen species

To investigate the generality of the preference for pathogenic substrates observed in our previous experiments, we tested whether flies also preferred substrates contaminated with other pathogens. To this end, we repeated the avoidance assays with three Gram-negative bacterial pathogens (*P. entomophila*; *P. aeruginosa* −PA14; *Providencia rettgeri)* and with the Gram-positive *Enterococcus faecalis*. Instead of repeating these assays on the set of 122 lines, we instead employed an advanced outcrossed population derived from the DGRP panel called the Ashworth Outcrossed Population (AOx Population). Briefly, this outbred population was established from 100 pairwise crosses of 113 DGRP lines in order to obtain an outbred population, with similar levels of genetic diversity present within the DGRP [49,53], which at the time of the experiment had been maintained for at least 90 generations of outcrossing. This approach was useful, as it allowed us to test pathogen avoidance on fewer flies and showed a mean response that was comparable to that of the 122 DGRP lines (Figures 1A and 2). Further, while the results using the individual DGRP lines included only female flies, here we included males and females which allowed us to test if pathogen preference was a female-specific response or common in both sexes.

Overall, we observed similar outcomes to the initial DGRP choice assay, and both male and female flies showed a general preference for the pathogen-contaminated substrate, particularly when exposed to *P. entomophila* (preference index (PI) = −0.33± 0.06; test if mean is different from zero: t = −5.5022, df = 119, p = 2.186e-07), *P. rettgeri* (PI= −0.34±0.05; t = −7.1034, df = 168, p = 3.294e-11), and *P. aeruginosa* (PI = −0.16±0.05; t = −3.0766, df = 158, p = 0.0025)(**Figure 2**). We found no evidence for pathogen preference (or avoidance) when flies were exposed to *E. faecalis* (PI= −0.05±0.060; t = −0.925, df = 158, p = 0.3564). We did however, observe clear pathogen avoidance when we increased the concentration of the PA14 from OD1 to OD100 (PI= 0.61± 0.06; t = 10.756, df = 111, p- < 0.001). This demonstrates that there are conditions under which flies will find bacterial cultures unpalatable, but such high concentrations of bacteria are unlikely to be common in nature. Under more realistic conditions, where concentrations are that of an overnight bacterial culture, flies did not avoid bacterial pathogens and in some cases showed a strong preference for pathogen-contaminated substrates. We considered the possibility that the starvation period experienced by flies prior to the FlyPAD assay may have influenced their preference [42], but we found similar preference patterns when we compared two experiments using either a short (4-6-hour) starvation period or a longer (18-24-hour) starvation period (Fig S1).

**Figure 2.**
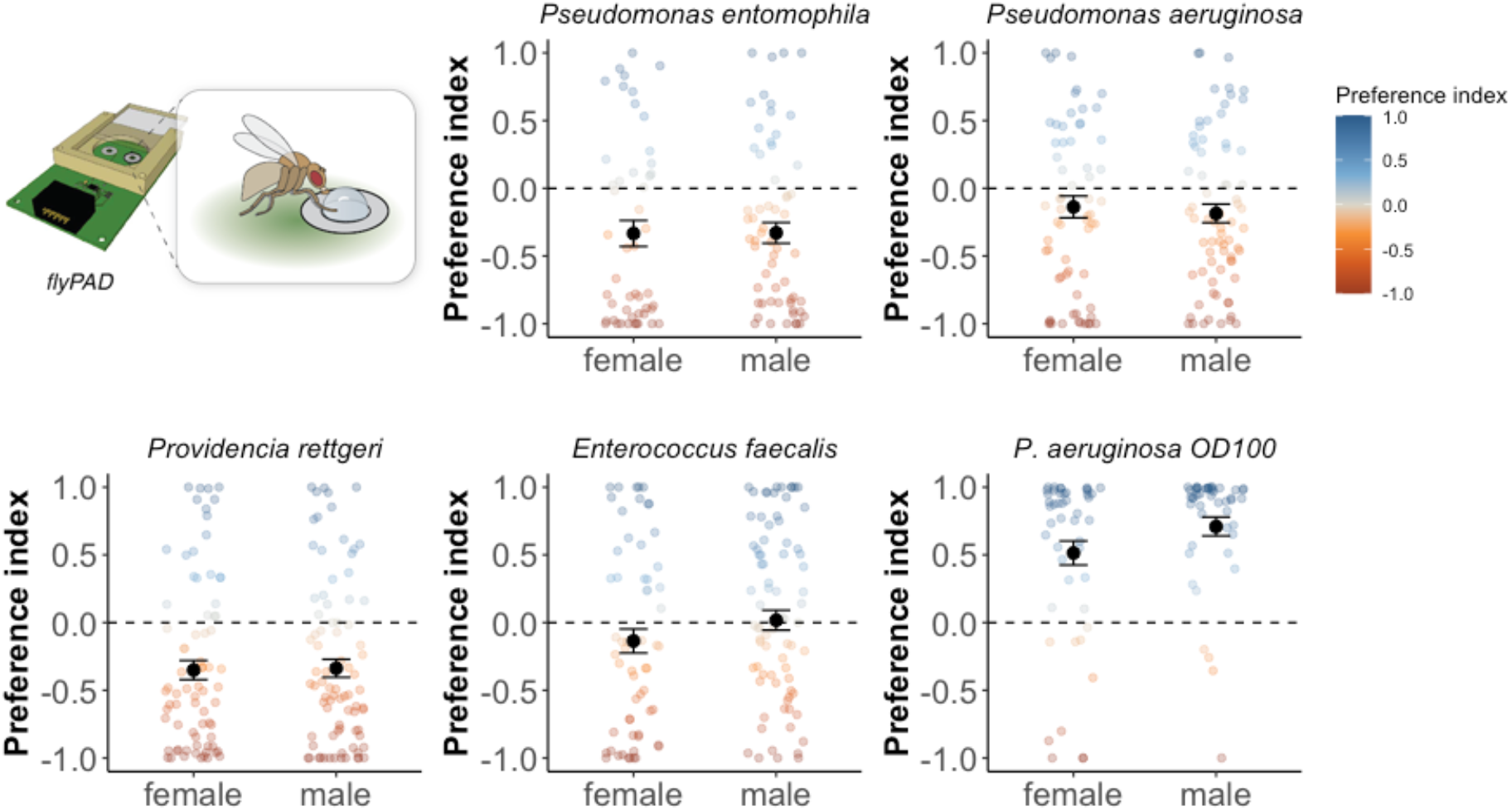
The preference index measured in response to different bacterial pathogens. Two-choice feeding assays were carried out as described for Figure 1 but using flies from an DGRP-derived advanced outbred population. We calculated the preference index (PI) in males and females given a choice of a clean substrate (5% sucrose) or OD_600_=1 *P. entomophila* (n= 55 females; 65 males), *P. aeruginosa* (n= 78 females; 81 males), *Providencia rettgeri* (n= 83 females; 86 males), *Enterococcus faecalis* (n= 73 females; 86 males), *or P. aeruginosa* (OD600=100; n= 56 females; 56 males). Each data point is the PI of an individual fly. Points are coloured according to the PI (blue indicates pathogen avoidance and red indicates pathogen preference). Black dots and error bars are the mean±SE of each group. FlyPAD image from https://flypad.rocks/.

### Variation in pathogen preference index is not driven by differences in susceptibility to bacterial infection

We hypothesized that the strong preference for pathogenic substrates could be driven by differences in susceptibility between fly lines, whereby lines with a strong preference for pathogenic substrates might also be less susceptible to bacterial infection. We tested this hypothesis in two ways. First, we chose the fly lines with either the highest or lowest preference index (ten lines per extreme) and we exposed these flies to an oral *P. entomophila* infection (OD_600_= 50) as an assay of infection susceptibility[54]. Most flies died within two days of infection and almost all flies had died within 96 hours. We could not distinguish between flies showing weak or strong preference for the pathogen substrate based on their mortality profiles (hazard ratio: 1.099± 0.07 SE; z=1.34, p=0.18), suggesting that lines with a very high attraction to pathogen substrates when feeding did not have greater capacity to resist or clear a bacterial infection (**Figure 3A**).

**Figure 3.**
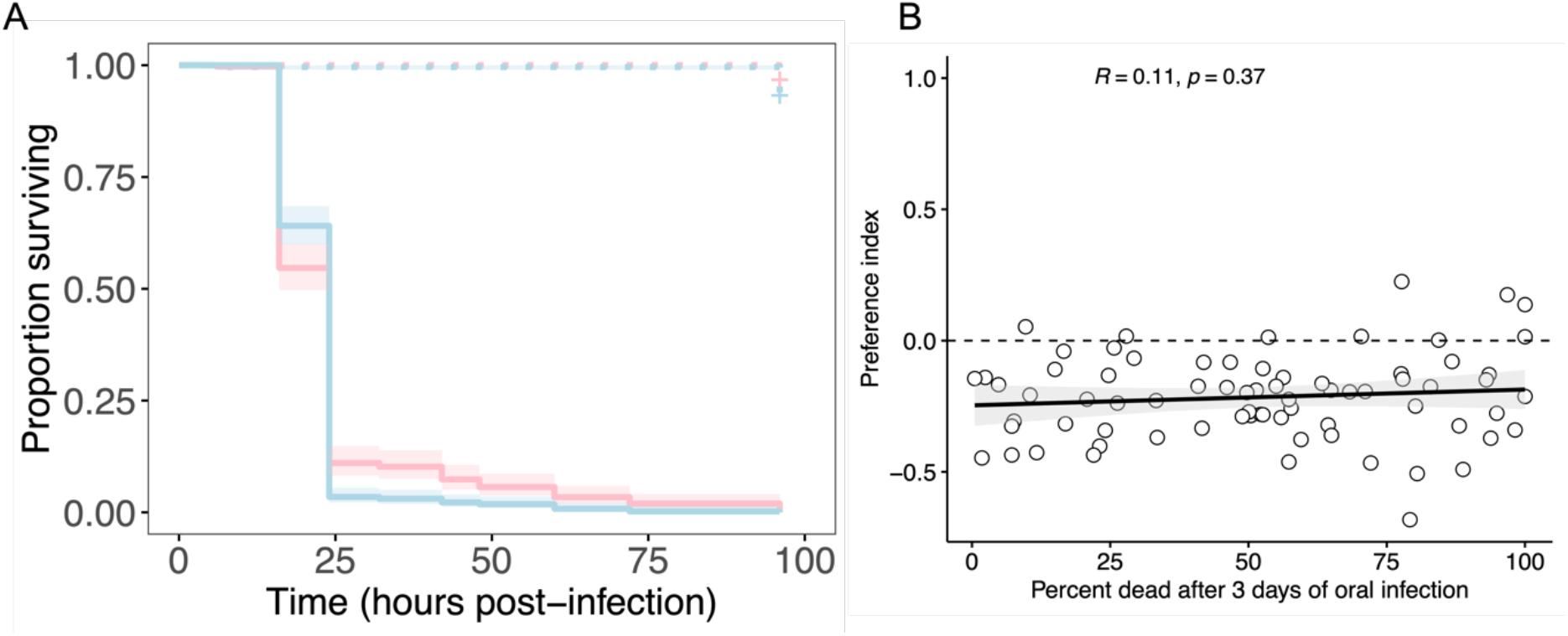
Testing associations between the preference index and infection susceptibility. A) Five lines with the highest (red) and five lines with the lowest preference index (blue) were challenged with an oral *P. entomophila* infection (as described in [54,55]) at a concentration of OD_600_ = 50; n = 7 replicate groups, with 10 flies per group (70 flies) per treatment. Crosses indicate censored flies that remained alive at 96h post infection. B) The relationship between the preference index and the percentage of flies that died after 3 days of oral infection with *P. entomophila*. Each data point is the line mean of preference measured for each DGRP line in Figure 1 and the survival of the same line as measured in [30]. R is the Pearson correlation coefficient, and the line shows the linear relationship, shaded by the 95% CI.

We went further to test the relationship between the measured preference index and the susceptibility of flies to an orally acquired, enteric *P. entomophila* infection. Here, we were able to use previously published data on susceptibility to *P. entomophila* oral infection in the DGRP, which contained 73 DGRP lines in common to our experiment [30]. In concordance with the oral infection of the lines showing extreme preference (**Figure 3A**), we found no significant association between the line means for the pathogen preference we measured, and the line means for survival of flies three days following oral infection (**Figure 3B**). Taken together, these analyses of infection data suggest that variation between fly lines in the preference for pathogenic substrates is unlikely to be driven by differences in susceptibility.

### A functional Immune Deficiency (IMD) pathway is not required for pathogen attraction

Both infection assays described above were designed to guarantee that flies acquire an infection, and therefore reflect susceptibility once the pathogen has established infection. However, we might expect the main immune-related driver (if any) of pathogen avoidance would be the susceptibility to acquire infection via the oral infection route, which may be under potentially distinct immune control. To further investigate a potential role of immunity in shaping the observed preference for pathogenic substrates, we took a functional genetic approach and performed the same two-choice assay using transgenic Drosophila lines with disrupted IMD-pathway function, which is the primary immune pathway involved in the response to gram-negative bacterial infection in Drosophila [56]. These experiments were also motivated by previous work showing that the IMD pathway, particularly the peptidoglycan receptors PGRP-LC and PGRP-LB, play a direct role in the detection of pathogen cues and subsequent neuroimmune signalling involved in pathogen avoidance behaviours [7–9].

We found that the control line *w*^*1118*^ displayed a preference for the pathogenic substrate (*P. aeruginosa* in this case), as observed in the DGRP panel, and the DGRP-derived AOx population. Disrupting the function of *Relish*, the main NF-kB transcription factor in the IMD pathway (**Figure 4A**), or the peptidoglycan receptors *PGRP-LB or PGRP-LC* did not significantly alter this preference (Figure 4), suggesting that IMD-mediated immunity is not involved in the observed preference for pathogenic substrates.

**Figure 4.**
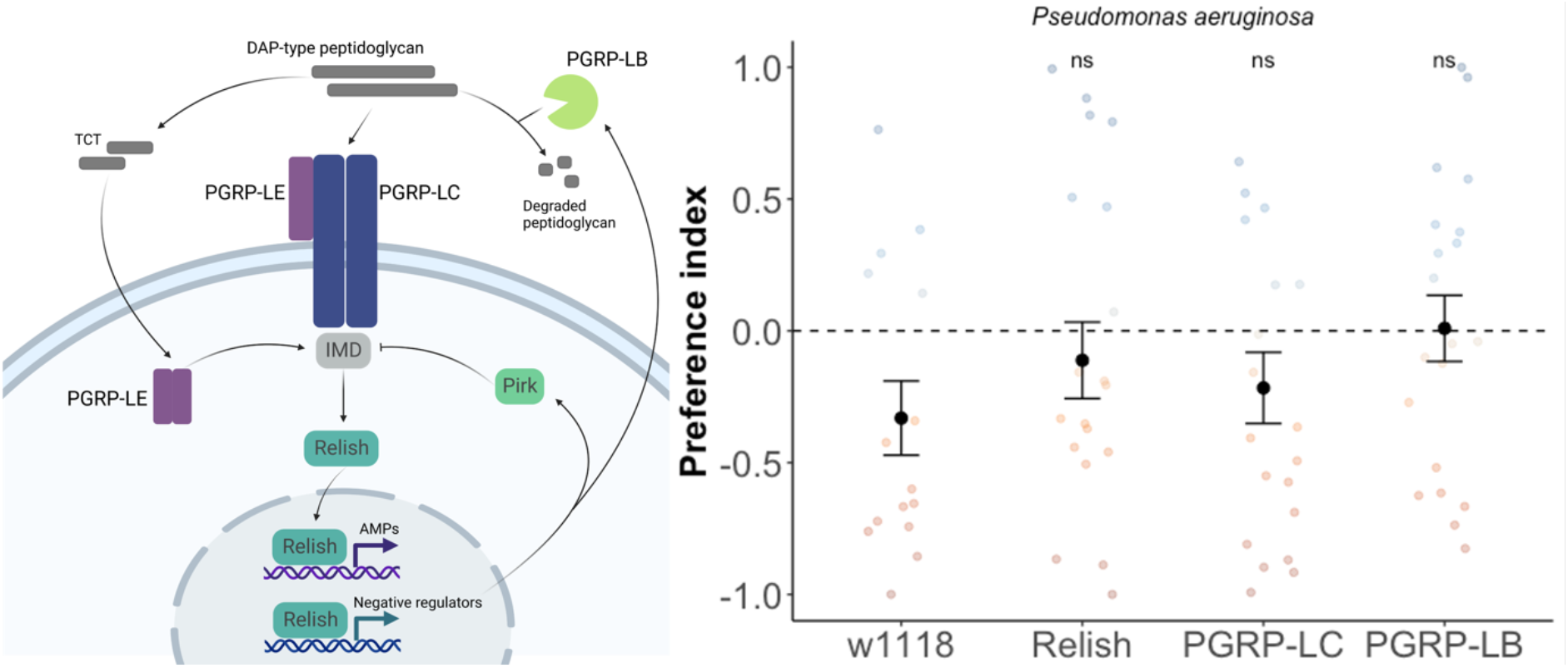
Pathogen preference measured in IMD loss-of-function lines. We used three transgenic lines with loss-of-function of specific IMD-pathway signalling components. Each line was isogenised onto *w*^*1118*^, which acts as a control with complete IMD signalling. Preference assays were carried out as described for experiments above (Fig 1 and Fig 2), negative values indicate a preference for the bacterial substrate. ns – non-significant difference tested using pairwise contrasts between *w*^*1118*^ and each loss-of-function line.

### Genotype-phenotype associations reveal novel genetic variants associated with pathogen preference

The DGRP is often employed as a powerful resource for identifying genetic variants underlying complex traits through genome-wide association studies (GWAS)[49,57,58]. Despite the low broad-sense heritability (Table 1) and that the 122 lines used in this study are likely unpowered to detect small effect variants [49], we decided to run a GWAS analysis in an attempt to identify any putative large-effect candidate variants associated with the phenotypic variation in the feeding preference.

GWAS analysis revealed a clear peak on chromosome 3R showing a strong association (p < 10^−8^) between variation in the preference index and several SNPs (Figure 5). To refine our analysis to the most promising candidate variants, we focused on variants with a p-value < 10^-6^, which resulted a list of 48 SNPs (**Table S1**). Notably, the top ten associations ranked by p-value all mapped to a gene-dense region spanning approximately 2000 bp on chromosome 3R. SNPs in this region mainly caused synonymous changes in three adjacent genes: *CG2321* and *CG2006* are coding proteins with nuclear expression but which are otherwise poorly described and have currently unknown functions. One of the SNPs in CG2006 also maps to *spase12*, which is essential for cell differentiation and development in Drosophila [59]. These genes are directly downstream of *protein tyrosine phosphatase 99A (ptp99A*), also included in the top 48 SNPs (Table S1). *ptp99A* has previously described functions in axon growth and guidance during neuronal development [60]. *ptp99A* has also been described as having a role in the regulation of food intake in a GWAS of Drosophila feeding behaviour[52]. Despite resulting in synonymous changes, all variants in this region of chromosome 3R were associated with a decrease in preference index, that is, attraction to the pathogen substrate (Figure 5).

**Figure 5.**
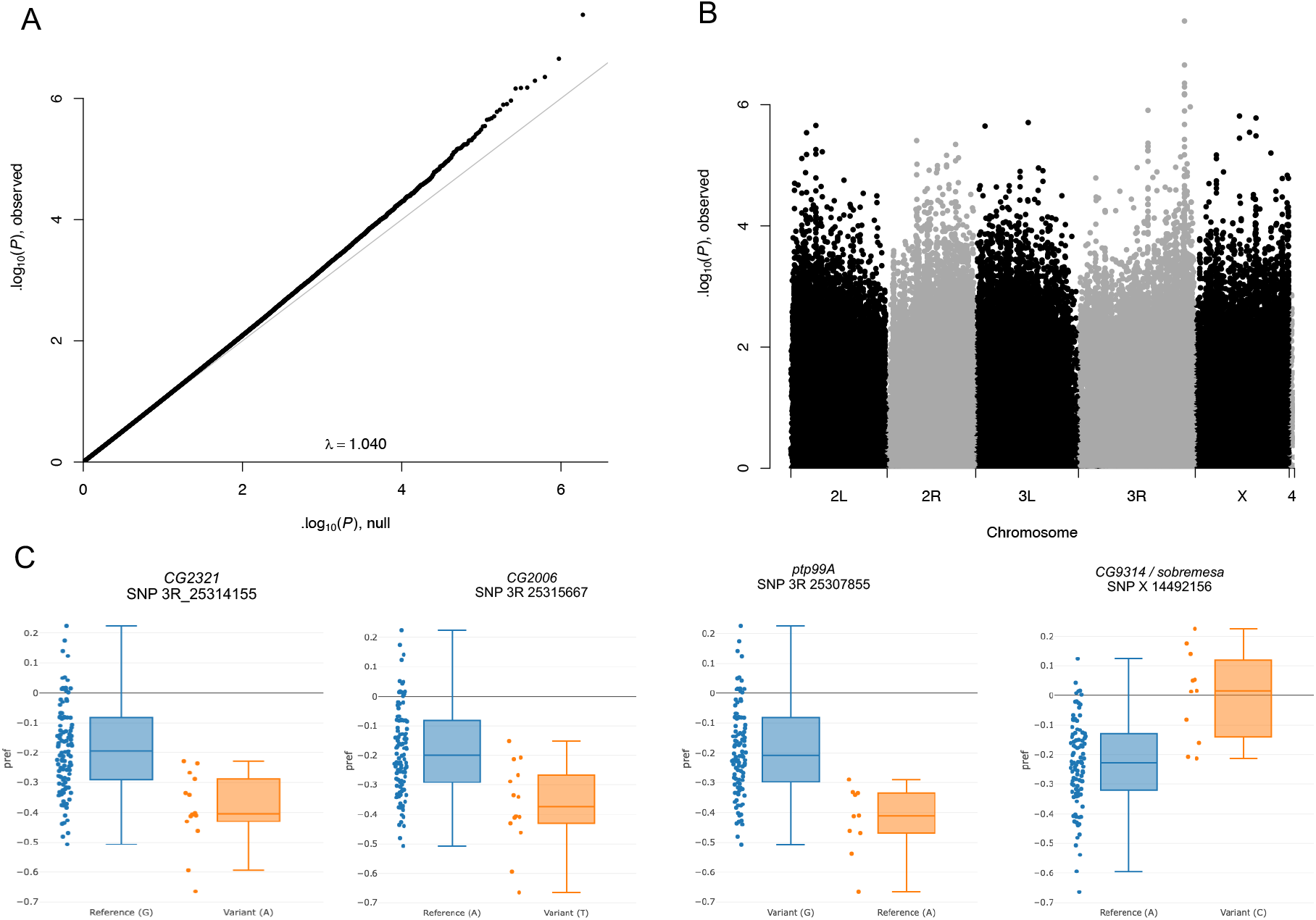
GWAS. A) The QQ plot shows quantile distribution of observed variant-phenotype association p-values compared to the distribution of expected p-values by chance; deviations above the diagonal line (and λ >1.00) indicate associations with a smaller p-value (more significant) than would be expected by chance. B) Manhattan plot showing single-SNP GWAS, testing the statistical association between the preference index and the SNPs in the genomes of the 122 lines tested. Each point is a SNP found along the genome, and fly chromosomes are labelled on the x-axis; the y-axis shows the −log10 p-values, where higher points indicate stronger associations. C) The preference index of DGRP lines carrying either the reference or variant alleles for the most significant SNPs on three genes clustered on chromosome 3R – *CG2321, CG2006, ptp99A*. We also plot *CG9314/ sobremesa* as the most significant variant associated with an increased preference index (greater pathogen avoidance). See Table S1 for a list of 48 SNPs with p-values <10^−6^.

Variants associated with increased avoidance of bacteria (a positive preference index) were much less common, though this is to be expected given the predominance of DGRP lines showing bacterial preference. However, within the top 48 variants, two independent SNPs were associated with increased avoidance, both located on the X chromosome and specifically on gene *CG9413*, also known as *sobremesa* (Figure 5). *sobremesa* encodes an SLC7 transmembrane amino acid transporter, which plays a crucial role in regulating amino acid homeostasis and feeding into signalling pathways that control developmental timing, growth, and feeding [61,62].s*obremesa* is expressed in larval glial cells and has been shown to modulate brain development in larval stage *D. melanogaster* [63,64]. While the function of *sobremesa* has only recently been described to be involved in the regulation of larval growth, it is also expressed in adult head and gut tissues (Flybase FBgn0030574; FB2024_02; [65]), so it is plausible that it functions as an amino-acid sensor in adult flies and could therefore modulate behavioural changes in response to their amino-acid environment.

### Potential fitness benefits of pathogen preference

Given the lack of association with immunity, and that the GWAS analysis indicated two different genes (*ptp99A* and *sobremesa*) with putative roles in feeding [52] or nutrient sensing[63], one possibility is that the preference for pathogen-contaminated substrates we observe may in fact be driven by a preference for amino-acids. Due the make-up of bacterial cell-wall components such as peptidoglycans, amino-acids were more abundant in the substrate containing the bacterial inoculum relative to the sucrose-only substrate. For example, the peptidoglycans of *Pseudomonas aeruginosa* are typically composed of an amino-acid chain including L-alanine, D-glutamic acid, Diaminopimelic acid (DAP), and D-alanine [66]. A plausible hypothesis is that by choosing a bacteria-contaminated substrate, flies are in fact seeking out higher protein content and that this diet-driven choice is presumably adaptive.

A strength of DGRP as a community resource is that many phenotypes have been carefully measured in the same lines in multiple labs [49,57,58]. We therefore analysed data from a previous study which had investigated the macronutrient tolerance in the DGRP by measuring the percentage of larvae that pupated and eventually emerged as adult flies under six different diets [67]. This work included a <u>H</u>igh <u>S</u>ugar <u>D</u>iet (HSD), and a <u>H</u>igh <u>P</u>rotein <u>D</u>iet (HPD), that are comparable to our 5% sucrose agar (clean) substrate and our pathogen substrate, respectively. This data shows that both aspects of fly developmental viability are generally higher in the HPD when compared to the HSD, regardless of the preference index we measured (Figure 6). This is broadly in agreement with our observation that almost all DGRP lines showed a preference for pathogen contaminated food which was relatively richer in protein (Figure 1A).

**Figure 6.**
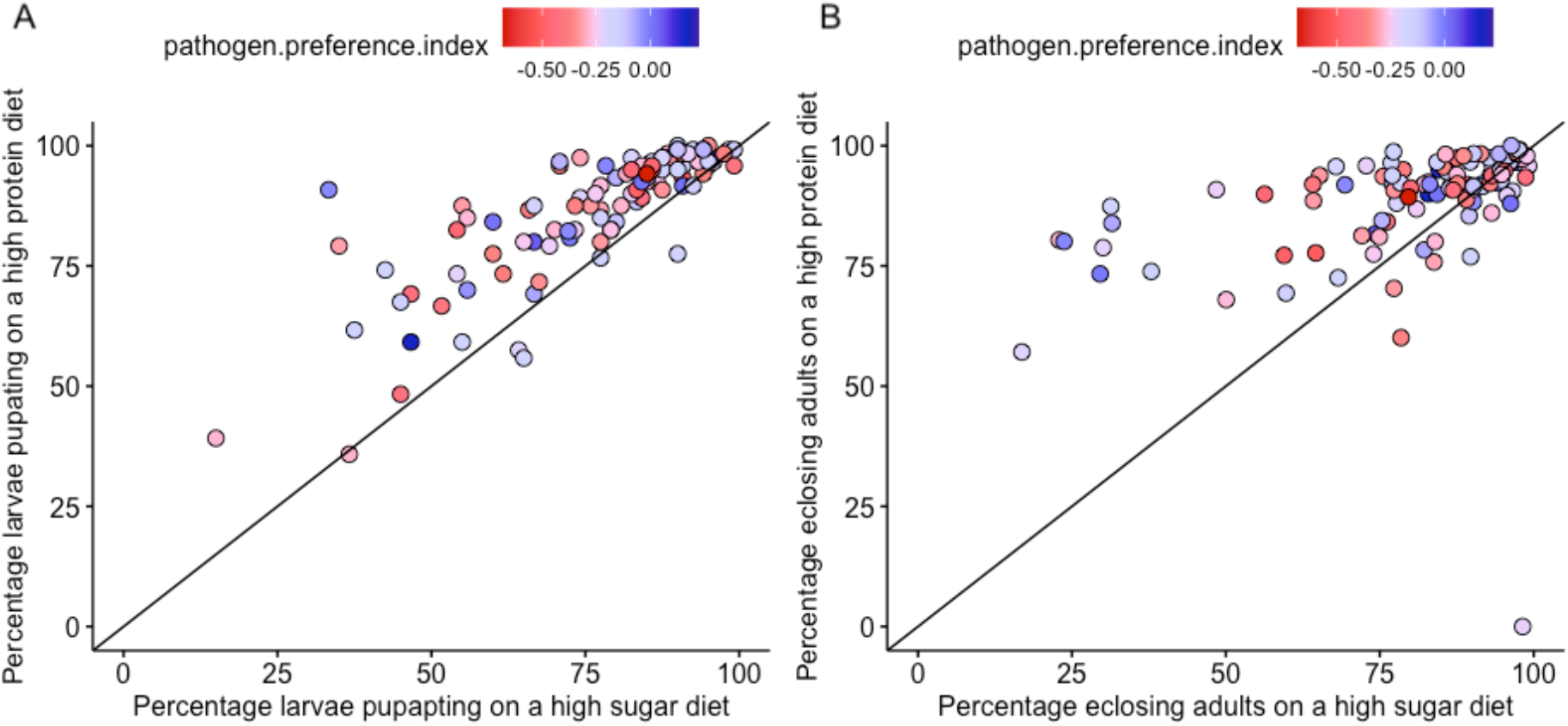
The effects of diet on Drosophila developmental traits. A) The percentage of larvae pupating and B) the percentage of eclosing adults, each on high protein vs a high sugar diet. Each data point is the line mean of a single DGPR line for each trait/diet. Data from [67].

The colour of each point indicates the preference index for each DGRP line as shown in Figure 1, where red indicates pathogen preference and blue indicates pathogen avoidance. The diagonal line indicated a correlation of 1 if traits are equal on both diets. Points above the diagonal indicate a higher trait value on the high protein diet.

## Discussion

Avoidance behaviours are often assumed to be adaptive, but it is notable how we know so little about the basic requisites for this assumption[2]. To be adaptive, avoidance must be amenable to respond to selection, which requires heritable phenotypic variation that this variation is associated with differences in fitness [50,68]. Yet, while elegant functional neurogenetic experiments have dissected the mechanisms underlying avoidance behaviours, especially in invertebrates [9], we lack measurements of phenotypic and genotypic variation in avoidance behaviours is most species [2,17,69]. Taking our results together, we find substantial intrapopulation variation in the likelihood to avoid infection, measured across 122 genetic backgrounds, although we find that this variation is only weakly heritable. Contrary to our initial expectation however, we found that most genotypes did not avoid the bacterial substrate, but instead preferred to feed on it.

We also hypothesised that behavioural avoidance would be associated with immune-mediated susceptibility. First, we might expect an avoidance-resistance trade-off where lines that are strongly attracted to pathogenic substrates may show higher resistance to infection as found previously in salmon [70] and sheep [71]. However, we did not find any evidence for trade-offs. Further, previous work has implicated specific components of the IMD signalling pathway, such as PGRPs, in the detections and signalling of bacteria-derived cues [7,8]. However, we did not detect any substantial effect of disrupting immune signalling on the preference for pathogen substrates. Instead, we argue that this preference is most likely driven by a dietary preference for protein. This is supported by the outcome of the genotype-phenotype association analysis, which implicated the role of *ptp99A*, previously linked to variation in food intake and *sobremesa*, an amino acid transporter, and also by the general fitness advantage of high protein diets compared a high sugar diet.

A potential limitation of our experimental design is the precise choice flies were presented with, which explicitly tested a preference between clean, sugar-only substrate, and an identical substrate containing an overnight culture of bacteria. This design does not allow distinguishing between a fly preference for consuming bacteria, or a bacteria-induced attraction. For example, some bacterial volatiles like ammonia and amines may attract flies, while short-chain fatty acids derived from bacteria also induce attraction [44–47]. However, while previous work has shown that flies detect and avoid odours associated with pathogenic infection [11], other work has found, as we have here, that flies initially prefer the odour of pathogenic bacteria - even when given a choice of two strains that are identical except for the production of a virulence factor [15]. While this bacterial preference was reduced if flies were fed with bacteria before the choice assay, this was the case whether or not the fed bacteria were pathogenic, suggesting that the main driver of the change in preference was not a learned avoidance of pathogens, but more likely a feeding/satiety-related response[15]. These results are concordant with ours, indicating that flies are generally attracted to bacterial substrates and that these choices are likely driven by a fly’s dietary amino-acid requirements.

Independently of the precise mechanism driving the preference for bacterial pathogens, this pattern of variation is likely to have important implications for the ecology and evolution of infection. In the simplest sense, broad attraction to pathogen-contaminated substrates will increase the likelihood of contact between flies and infectious sources. Further, given that we find there is no association with susceptibility, this should translate into a higher prevalence of infection and its associated fitness costs. In support of this prediction, a model of behavioural resistance to parasites – which under some assumptions is equivalent to pathogen avoidance – shows that broadly, lower average levels of avoidance are expected to lead to higher infection prevalence [38]. Other work investigating the role of individual trait heterogeneity on the population-level epidemic outcomes found that populations with high levels of behavioural variance (related to avoidance and relevant for contact rates between infectious and non-infectious individuals) were more likely to experience more severe epidemics[24]. The evolutionary implications of higher infection prevalence, in turn, will depend on the relative costs of infection and of both behavioural and other physiological and immune defences [36,38].

In sum, avoidance behaviours no doubt play an important role in host defence by avoiding the detrimental effects of infection, but also by allowing individuals to avoid the physiological costs of immune deployment [28,72]. It is important to recognise, though, that individuals are likely to vary in their propensity to avoid sources of infection, particularly when these come at the cost of not feeding or reproducing. In some cases, what is perceived as avoidance may instead be the by-product of selection on these other fitness-enhancing activities [19,69]. Understanding the ecological and evolutionary drivers of avoidance, and how avoidance is likely to drive the ecology and evolution of hosts and pathogens therefore remains an important and understudied topic of research, and requires more widespread measurement of the extent to which individuals vary in avoidance and how these behavioural choices may impact fitness [2,16,26,73,74].

## Methods

### Fly lines and outbred population

We measured the choice index in individual, mated female flies from 122 lines from the Drosophila Genetic Reference Panel (DGRP), a panel of fully-sequenced fly lines originating from the same population, and therefore offers a useful snapshot of natural genetic variation in pathogen avoidance behaviour [49,58]. These specific lines have been maintained in our lab since 2013 and differ from the original DGRP in that they have been previously cleared of from the endosymbiont *Wolbachia* by rearing with the antibiotic Tetracycline for two generations [75]. Other lines used were *w*^*1118*^ (Vienna *Drosophila* Resource Center); *Rel*^*E20*^ (*relish - IMD* pathway regulator [76]); PGRP-LC (Peptidoglycan receptor protein-LC; BDSC_12500)[77]; PGRP-LB (BDSC_55715)[78]. All flies were raise in identical conditions (25ºC±1ºC; 12:12 light:dark cycle), and maintained at similar densities by placing 5 females and 2 males per vial for 48 hours, and using their progeny as experimental flies.

In some assays, we employed an outbred fly population. The Ashworth Advanced Outcrossed (AOx) population is a large, lab adapted, outbred *Drosophila melanogaster* population derived from the DGRP panel [58]. The AOx was originally established in October 2014 by setting up 100 crosses of 113 DGRP lines [79], and has been maintained since then as an outbred population on a 14-day generation cycle with census size of between 3000-4000 adults in each generation. Every generation, eggs are collected from a population cage (a plastic container: 30 cm × length × 20 cm width × 20 cm height) and dispensed into 8 oz bottles (57 mm diameter x 103 mm height containing cornmeal-sugar-yeast medium[80] at a density of 100 (±20) eggs per bottle. 25 bottles were set up at each generation and incubated at 25 °C, 12:12 hour LD cycle. The development time of these flies is 9-10 days. Two-to-three weeks post egg collection, all adults present in bottles are transferred to a population cage and provided with two fresh apple agar plates (in a 90 mm Petri dish) supplemented with ad libitum yeast paste in the centre of each plate. 6-8 hours later all eggs are collected by careful washing and distributed to new bottles to begin the next generation.

### Bacterial strains and culture conditions

Depending on the experiment we used the following bacterial strains: *Pseudomonas entomophila* [55,81], *P. aeruginosa* [82,83], *Providencia rettgeri* [84,85], Enterococcus faecalis [86]. Unless otherwise stated, all bacterial cultures were prepared in the same way. A single colony stored at −70ºC in 100µl of 25% glycerol was inoculated in 10mL Luria-Bertani (LB) Broth and left to grow overnight in an orbital shaker at 37°C, 140 rpm. Overnight cultures were adjusted to OD_600nm_=1 and mixed 1:1 with a 2% agar, 5% sucrose LB broth – see FlyPAD assay below). In experiments using higher bacterial concentrations, overnight cultures were grown until exponential growth phase was reached (OD_600nm_ 0.6-0.8). This culture was then divided equally into 50mL falcon tubes and centrifuged at 2500 rpm for 15 minutes to form pellets and the supernatant removed. The pellets were resuspended by shaking and recombined in a single falcon tube, which was centrifuged again. The supernatant was removed, and the pellet resuspended in the appropriate volume of LB broth to reach OD_600nm_=50 (in the experiment with immune deletion lines) or OD_600nm_=100 (for one of the *P. aeruginosa* feeding assays).

### FlyPAD choice assay

We used the FlyPAD system for tracking, recording, and analysing the feeding behaviour of fruit flies in real-time [42]. Two-to-six-day-old flies were wet-starved (using water-soaked cottonwool plugs) overnight for 18-22 hours prior to each assay. Following the starvation period, flies were placed individually within a FlyPAD arena previously prepared with two choice substrates, one on each electrode. The “clean” choice substrate contained (LB broth, 2% agar & 5% sucrose); the infectious substrate was identical, but the sucrose was replaced with 5% of an overnight bacterial culture, adjusted to OD_600_=1. Each FlyPAD arena received 1ul of each substrate, carefully pipetted onto each electrode, and choice assays lasted 30 min [42]. Each DGRP line was replicated between 16-24 times, in blocks of 32 flies. We carried our four blocks each day, all between 10:30-15:00, randomising the genotypes between blocks to minimise any potential time-of-day effects. The order in which the substrate (clean or bacterial) was pipetted on to the FlyPAD was also alternated between blocks to account for any potential pipetting bias. We measured pathogen avoidance as a preference index, calculated over 30 minutes, as *(sips*_*clean*_ *-sips*_*pathogen*_*) / sips* _*total*_, that is, the difference in sips taken by an individual fly on either pathogen-contaminated substrate relative or a clean alternative, relative to the total number of sips. Positive values of the preference index are suggestive of pathogen avoidance, where a value of 1 indicates complete preference for the clean substrate; negative values suggest a pathogen preference, where −1 indicates complete preference for the pathogen-contaminated substrate; 0 indicates an equal number of sips on each substrate.

### Experimental infection of extreme preference lines

An oral infection experiment was carried out in order to establish whether those DGRP lines which showed the highest and lowest preference for *P*.*entomophila*-contaminated food varied in their susceptibility to infection. The 10 DGRP lines showing the highest and lowest preference for the *P*.*entomophila*-contaminated food were selected. Flies were reared and treated as above prior to experimentation. For each line, 7 x cohorts of 10 flies were fed a bacterial infected food source and 5 x cohorts of 10 were fed an alternative control food source. Prior to oral exposure, flies were wet-starved in cohorts of 10 for ∼6-8 hours before being added to the infection vial or corresponding control vial. Infection vials consisted of 7 mL Bijou tubes with 500 µL 8% sugar-agar added to the inside of the lid. The sugar-agar was left to set and topped with a filter disc, onto which 100 µL of *P*.*entomophila* suspended in 5% sucrose solution at O.D_600_ 50 was pipetted. Control tubes were set up in the same manner but 100 µL of 5% sucrose solution used instead. The bacterial solution and control solution were left to soak into the filter disc for 20 mins before flies were added. Flies were left in infection tubes for 24 hours and then moved to standard vials for survival monitoring. To confirm the infection status of experimental and control flies a subset of flies from each line were surface sterilized in 70% ethanol, followed by 3xdistilled H_2_O before being homogenised in 1xPBS. Both the 3xdistilled H_2_O wash-offs and homogenised flies were plated on LB agar. Plates were incubated for 24 hours at 30 °C before infection status was confirmed; homogenised flies orally exposed to *P*.*entomphila* were infected with the bacterium and those from control tubes were not.

### Phenotype-Genotype Association Analysis

We carried out Phenotype-Genotype Association Analysis using well established analysis pipelines[49,58]. Specifically, we used the GWAS analysis as implemented by DGRPool (https://dgrpool.epfl.ch/ [57], which uses Plink2 v2.00a3LM (1 July 2021) and runs all variants in the DGRP database, using the dm3 genome assembly (4,438,427 variants: 3,963,420 SNPs, 293,363 deletions, 169,053 insertions and 12,591 MNPs). While the DGRP lines used in this study have been previously cleared of the endosymbiont *Wolbachia* (see above), the DGRPool implementation of GWAS includes the *Wolbachia* status of the lines and five major insertions as covariates in the model[87]. We evaluated single marker associations using preference index line means with common variants and then used the resulting quantile-quantile QQ plots and Manhattan plots to identify strong associations between genomic variants and variation in each phenotype. We focused on candidate variants with p < 10^−6^ that mapped to coding genes or that that occurred within 5Kb of a coding gene and a minor allele frequency (MAF) >5% [57].

### Data analysis

All data and analysis code is available at https://doi.org/10.5281/zenodo.11149149 [88]. All data was analysed in R (v. 4.2.2) and RStudio (2023.12.1+402), using *tidyverse* for data manipulation and *ggplot2* to plot all figures[89]. We used the package *lme4* [90] to fit generalized linear mixed models (GLMM-using the function “*glmer*”) and linear mixed models using the function “*lmer*”; the package *lmerTest* was used to extract *p*-values from models[91]. We used the package *performance* [92], using the “*check_model”* function to evaluate the model fits, to test for severe deviations from gaussian-distributed residuals, and to test for overdispersion. To analyse the preference index among 122 DGRP lines, we fitted a constant (intercept-only) linear mixed model to predict ‘*preference*.*index’*. The model included *line, day*, and *replicate* (individual fly) as random effects:

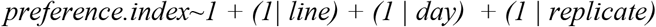

Variance components were extracted using the *VarCorr* package, and broad-sense heritability was calculated as the genetic variance *Vg* (the variance explained by DGRP line) divided by the total phenotypic variance *Vp* (the residual model variance), where *Vp = Vg + Ve* (the environmental variance)[49,50]. We used the package *coxme* to fit cox proportional hazards models with random effects [93]:

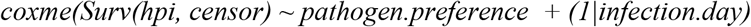

Differences between lines with a low or high preference for pathogen substrates were inferred from the significance of *exp(coef)* from the coxme summary, which provide the hazard ratio. Correlations between traits were carried out on the line means for each trait by testing the significance of the Pearson correlation coefficient.

## Acknowledgments

We thank Angela Reid and Alison Fulton for help with media preparation. We acknowledge funding and support from the Branco Weiss fellowship and a Chancellor’s Fellowship to PFV. PT was funded by a University of Edinburgh scholarship, through the Wellcome Trust PhD Programme in Hosts, Pathogens and Global Health awarded to Keith Matthews (grant 218492/Z/19/Z). For the purpose of open access, the author has applied a Creative Commons Attribution (CC BY) license to any Author Accepted Manuscript version arising from this submission.

## Author Contributions

KMM and PFV conceived and designed the experiments. KMM and PT performed the experiments. PFV analysed the data. PFV wrote the paper with additional inputs from KMM and PT.

## Data accessibility statement

All data and analysis code is available at https://doi.org/10.5281/zenodo.11149149 [88]. An earlier preprint is available from the biological preprint server *bioRxiv* at https://www.biorxiv.org/content/10.1101/2024.05.09.593162v1[94].

## Competing Interest Statement

The authors declare no competing interests

## Notes

https://doi.org/10.5281/zenodo.11149149

### Competing Interest Statement

The authors have declared no competing interest.

### Summary of Updates

Some inaccuracies were corrected throughout, especially to clarify details of the methodology. A section explaining experimental infections was added.

