## Supplementary Material for "Genetic variation in trophic avoidance shows fruit flies are generally attracted to bacterial pathogens"

| **#CHROM** | **POS** | **Gene** | **SNP** | **REF** | **ALT** | **BETA** | **P** |
| --- | --- | --- | --- | --- | --- | --- | --- |
| 3R | 25314857 | CG2321 | SYNONYMOUS_CODING | A | G | -0.115829 | 4.17E-08 |
| 3R | 25313819 | CG2321 | SYNONYMOUS_CODING | C | T | -0.123167 | 2.21E-07 |
| 3R | 25315632 | CG2006 | SYNONYMOUS_CODING | G | T | -0.115929 | 4.43E-07 |
| 3R | 25314155 | CG2321 | SYNONYMOUS_CODING | G | A | -0.114894 | 5.10E-07 |
| 3R | 25313060 | CG2321 | SYNONYMOUS_CODING | G | A | -0.094917 | 6.63E-07 |
| 3R | 25307855 | Ptp99A | SYNONYMOUS_CODING | G | A | -0.134724 | 6.72E-07 |
| 3R | 25313591 | CG2321 | SYNONYMOUS_CODING | C | A | -0.107413 | 6.87E-07 |
| 3R | 26691619 |  |  | G | A | -0.094659 | 1.09E-06 |
| 3R | 16598951 | Syp | INTRON | C | A | -0.142609 | 1.24E-06 |
| 3R | 25315560 | CG2006 | SYNONYMOUS_CODING | C | T | -0.100225 | 1.27E-06 |
| X | 10545444 | X11Lbeta | INTRON | C | G | -0.079119 | 1.54E-06 |
| X | 14492348 | CG9413 | INTRON | C | T | 0.112311 | 1.66E-06 |
| 3L | 12533325 | sowah | INTRON | G | A | -0.093392 | 1.97E-06 |
| 3R | 25313528 | CG2321 | SYNONYMOUS_CODING | A | G | -0.10193 | 2.12E-06 |
| 2L | 5964533 | psd | INTRON | A | T | -0.110105 | 2.21E-06 |
| 3L | 2224337 |  |  | CAGTAA | G | -0.121324 | 2.26E-06 |
| X | 12951840 | rad | INTRON | T | G | -0.089616 | 2.86E-06 |
| 2L | 3741950 | CG10019 | INTRON | C | A | -0.079468 | 2.90E-06 |
| X | 14492156 | CG9413 | INTRON | A | C | 0.125423 | 3.27E-06 |
| X | 10545363 | X11Lbeta | INTRON | C | T | -0.075655 | 3.57E-06 |
| 3R | 25308025 | Ptp99A | SYNONYMOUS_CODING | G | A | -0.106788 | 3.75E-06 |
| 2R | 7043676 | CG13229 | INTRON | T | C | -0.071153 | 3.91E-06 |
| 3R | 16597348 | Takl1 | INTRON | A | C | -0.122411 | 4.31E-06 |
| 2R | 16339680 | CG12484 | INTRON | C | CG | -0.086552 | 4.53E-06 |
| 3R | 16597398 | Takl1 | INTRON | G | ATACAAATTGTAAC | -0.116632 | 4.88E-06 |
| 3R | 16597400 | Takl1 | INTRON | G | GC | -0.116632 | 4.88E-06 |
| 3R | 25315667 | CG2006 | NON_SYNONYMOUS_CODING | A | T | -0.104616 | 4.97E-06 |
| 2L | 5964532 | psd | INTRON | C | A | -0.104135 | 5.50E-06 |
| 3R | 16597350 | Syp | INTRON | C | A | -0.115708 | 5.74E-06 |
| 3R | 16597353 | Syp | INTRON | GCA | TT | -0.115708 | 5.74E-06 |
| 3R | 16597358 | Syp | INTRON | AC | T | -0.115708 | 5.74E-06 |
| 2L | 7501580 | RapGAP1 | INTRON | T | C | -0.097126 | 5.99E-06 |
| X | 18048621 | Frq1 | UPSTREAM | A | C | 0.083002 | 6.28E-06 |
| 2L | 5966575 | CG13992 | SYNONYMOUS_CODING | T | C | -0.074661 | 6.55E-06 |
| 2L | 3741963 | CG10019 | INTRON | G | C | -0.075795 | 6.64E-06 |
| 3R | 25313303 | CG2321 | SYNONYMOUS_CODING | C | A | -0.10107 | 6.72E-06 |
| X | 5038044 | rg | INTRON | C | CGCATTAA | -0.070703 | 6.76E-06 |
| 2R | 14186448 | sbb | INTRON | A | T | -0.089145 | 6.85E-06 |
| 3R | 24389930 |  |  | C | T | -0.130964 | 7.26E-06 |
| 2R | 17034635 | Hmg-2 | SYNONYMOUS_CODING | G | A | -0.108841 | 7.53E-06 |
| 2L | 2601775 | CR44789 | UPSTREAM | A | G | -0.099239 | 7.73E-06 |
| X | 5038045 | CG15465 | INTRON | ACTCTTTG | A | -0.074439 | 7.77E-06 |
| 2R | 16339686 | CG12484 | INTRON | GG | G | -0.085785 | 8.44E-06 |
| 3R | 16597403 | Syp | INTRON | C | G | -0.119897 | 8.62E-06 |
| 3R | 25937924 | sima | INTRON | A | G | 0.086047 | 9.46E-06 |
| 2R | 7113018 | CG7737 | SYNONYMOUS_CODING | T | C | 0.130366 | 9.61E-06 |
| 2R | 14186405 | CG18539 | UTR_3_PRIME | T | A | -0.085913 | 9.93E-06 |

**Table S1. Summary of GWAS** results, listing the top 48 variants, ranked by lowest to highest p-value. This lists only includes variants with p<10^-6^.

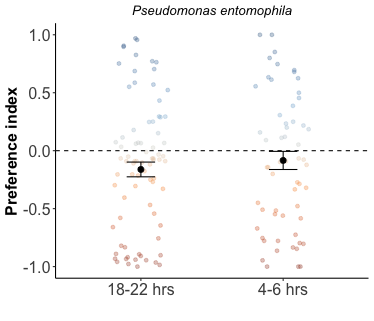

**Figure S1. Testing the effect of prior starvation period.** Prior to the FlyPAD choice assay, flies were starved for either overnight (18-22 hours n=83, 3–5-day-old female flies taken from an outbred population) or for a shorter starvation period of 4-6 hours (n=69 female flies from the same outbred population) on the day of the assay. No differences were detected in the average preference index.
